# NDC80 status pinpoints mitotic kinase inhibitors as emerging therapeutic options in clear cell renal cell carcinoma

**DOI:** 10.1101/2023.03.07.531634

**Authors:** Cheng Hu, Weiming Lin, Kemeng Zhao, Guiyou Tian, Xiangquan Kong, Guangcheng Luo, Dieter A. Wolf, Yabin Cheng

**Affiliations:** State Key Laboratory of Stress Biology and Fujian Provincial Key Laboratory of Innovative Drug Target Research, School of Pharmaceutical Sciences, Xiamen University, Xiamen, China; Department of Internal Medicine II, Klinikum rechts der Isar, Technical University Munich, Munich 81675, Germany; Ganzhou Key Laboratory for Drug Screening and Discovery, School of Geography and Environmental Engineering, Gannan Normal University, Ganzhou 341000, China; Department of Radiation Oncology, Xiamen Humanity Hospital, Fujian Medical University, Xiamen, China; Department of Urology, Zhongshan Hospital, Xiamen University, Xiamen, China; Westlake Laboratory of Life Sciences and Biomedicine and School of Life Sciences, Westlake University, Hangzhou 310024, China

## Abstract

∼30% of clear cell renal cell carcinoma (ccRCC) patients present with metastatic disease at the time of diagnosis, causing a dire 5-year survival rate of 13%. Whereas anti-PD-1 immunotherapy has improved survival, a strong need remains for new therapeutic options. Using integrated network analysis, we identified the mitotic regulator NDC80 as a predictor of ccRCC progression. Overexpression of NDC80 fosters the malignant phenotype by promoting cell cycle progression through S phase as well as boosting glycolysis and mitochondrial respiration. Despite high levels of immune infiltration, particularly derived from tumor resident CD8+ T cells with an exhausted phenotype, NDC80 defines a class of ccRCCs that poorly respond to immune checkpoint blockade. Instead, bioinformatics identified NDC80-high ccRCCs as sensitive to inhibitors of mitotic kinases, PLK1 and AURK, therapeutic approaches we validated in cell lines and mouse xenograft studies. Thus, NDC80 status pinpoints mitotic kinase inhibitors as promising therapeutic options in difficult-to-treat ccRCCs.

## INTRODUCTION

Clear cell renal cell carcinoma (ccRCC) is the most common subtype of renal cancer and accounts for up to 80% of all cases ^1^. Patients with localized ccRCC have survival rate of 70%-90% after partial or radical nephrectomy, but 5-year survival drops to 13% in the ∼20% of patients who progress to metastatic disease ^2^. ccRCC is typically insensitive to standard cytotoxic chemotherapy ^3^. Tyrosine kinase inhibitors targeting VEGF signaling and immune checkpoint inhibitors have improved ccRCC patient outcome ^4^. However, many patients do not or only initially respond to these treatments. The lack of clinically relevant predictive biomarkers severely hampers the therapeutic decision process, and their identification thus represents a priority in ccRCC ^5,6^. Similarly, novel therapeutic targets for ccRCC are urgently needed.

Metabolic reprogramming as a consequence of the inactivation of the von Hippel-Lindau tumor suppressor gene VHL is a near universal property of ccRCCs ^7^. VHL mutations induce angiogenesis and cell proliferation by elevating the expression of hypoxia-inducible factor 1 and 2 alpha (HIF1α and 2α) in ccRCC. HIF1α and HIF2α stimulate the expression of GLUT1 (SCL2A1), pyruvate dehydrogenase kinase (PDK1), and lactate dehydrogenase A (LDHA) thereby shifting glucose metabolism towards glycolysis. Besides, HIF1α also mediates mitochondrial autophagy via BNIP3 and BNIP3L ^8,9^. Therefore, metabolic pathway regulators might be promising biomarkers and/or drug targets in ccRCC.

Recently, NUF2 was identified as a factor that is overexpressed in ccRCC and correlates with poor patient prognosis ^10^. NUF2 is one of four subunits of the nuclear division cycle 80 (NDC80) complex which is a core component of the outer kinetochore and a mitotic factor implicated in cancer progression through its maintenance of genome stability and ploidy ^11,12^. Using orthogonal data mining approaches, we have identified in the present study another subunit of the NDC80 complex, the name giving NDC80 subunit (also called HEC1) as a prognostic marker in ccRCC. Overexpression of NDC80 was previously observed in a variety of cancers and implicated in tumorigenesis ^13–16^, making NDC80 a possible therapeutic target in breast cancer and colon cancer ^17,18^. However, the potential of NDC80 as a biomarker predicting therapeutic outcome in ccRCC has not been explored. Our study reveals NDC80 as a novel metabolism-related biomarker predicting the responsiveness of difficult-to-treat ccRCCs to mitotic kinase inhibitors thus pinpointing urgently sought new therapeutic options in this devastating disease.

## RESULTS

### Integrated network analysis implicates NDC80 in ccRCC progression

To decipher cellular pathways involved in ccRCC and identify potential disease biomarkers and drug target pathways, mRNA expression data from four GEO cohorts were obtained for 182 ccRCC samples and 127 normal kidney tissue samples. As outlined in **Figure S1**, differentially expressed genes (DEGs) were identified in the individual datasets (**Figure S2A-2D**) and subjected to comprehensive bioinformatic analysis. We focused on the 338 upregulated and the 405 genes common to all four cohorts (**Figure S2E-S2F**). As expected, these lists of DEGs were enriched for Gene Ontology (GO) terms related to kidney development/function and hypoxia (**Figure S2G**) as well as the KEGG pathways HIF1 signaling and glycolysis/gluconeogenesis among others (**Figure S2H**).

To further refine the functional pathway analysis toward potential drug targets, a protein-protein interaction (PPI) network of DEGs was assembled from the STRING database. Clustering revealed multiple densely interconnected subnetworks that were enriched for distinct Reactome pathways, including central carbon metabolism (glycolysis, TCA cycle, oxidative phosphorylation), immune regulation (T cell activation, TLR signaling), mitotic control (PLK1 and aurora kinase signaling), and HIF1/2a transcriptional activity (**Figure 1A**). These enrichments are consistent with the hallmark loss of VHL resulting in HIF1 activation and metabolic reprogramming frequently observed in ccRCC ^7^. The data also indicate changes in immune signaling and S/M cell cycle control.

**Figure 1.**
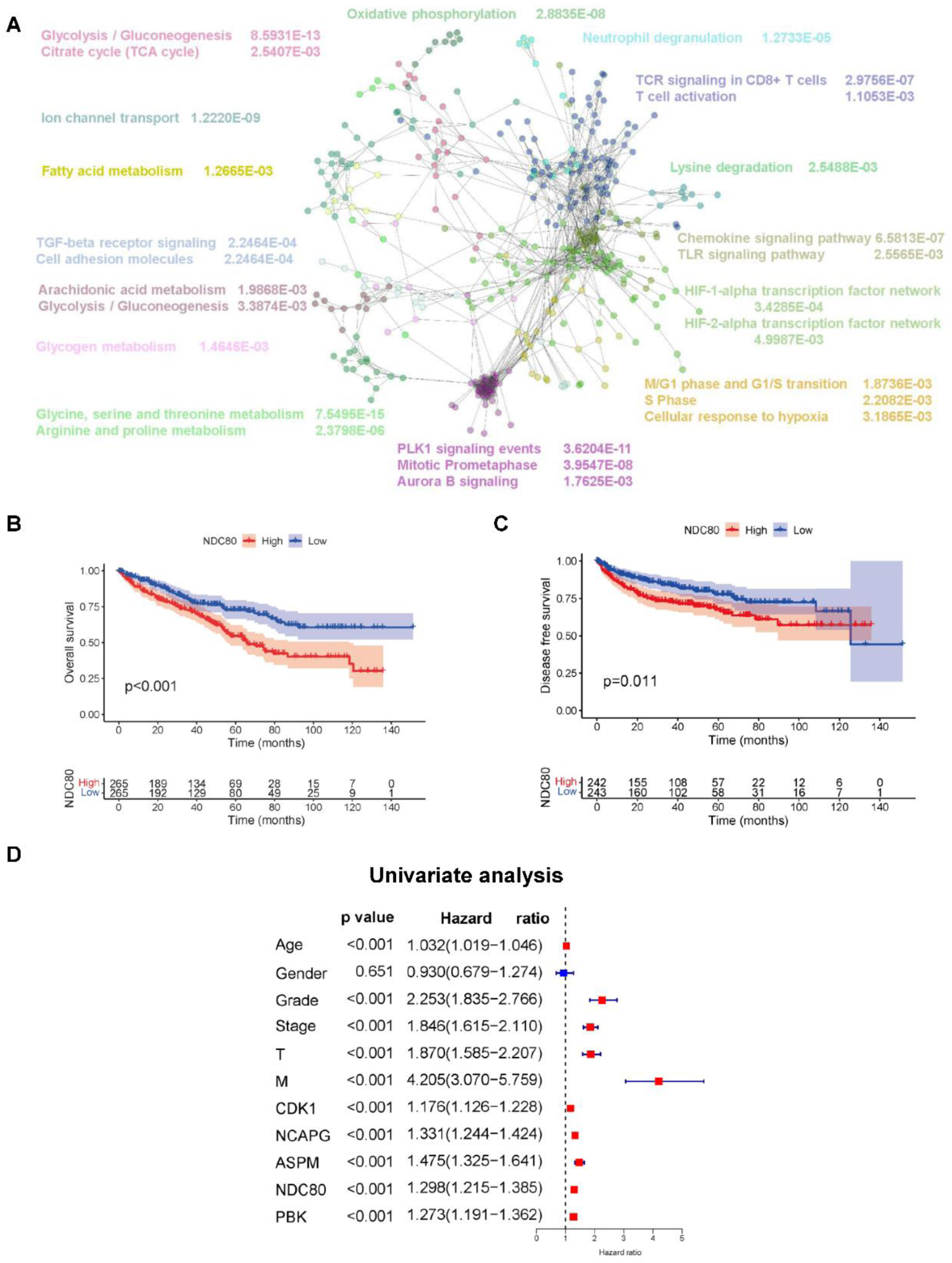
Integrated network analysis implicating NDC80 as a prognostic factor in ccRCC. (**A**) Reactome pathways analysis of a protein-protein interaction network constructed from genes differentially expressed in ccRCC according to four GEO datasets. (**B, C**) Kaplan–Meier plots of overall survival (OS) and diseases free survival (DFS) of ccRCC patients based on the expression of NDC80 in the TCGA database. Patients were divided into NDC80-high and NDC80-low groups based on the median NDC80 mRNA expression level. (**D**) Univariable analysis for overall survival (OS) of ccRCC patients. P values and hazard ratios including 95% CI (confidence interval) are showed.

Focusing on the most densely interconnected mitotic control network, the Cyhubba app for Cytoscape was used to identify hub proteins using the Maximal Clique Centrality (MCC) method. ASPM, CDK1, NCAPG, NDC80, CCNA2, DLGAP5, NUSAP1, PBK, KIF11, TTK were identified as the top 10 hub proteins (**Figure S3A**). Using quantitative LC-MS data from the Clinical Proteomics Tumor Analysis Consortium (CPTAC, ^19^) through the UALCAN web portal ^20^, we found that the expression of 5 of the 7 top hub proteins detected by LC-MS (CDK1, NCAPG, ASPM, NDC80, PBK) was increased in ccRCC (**Figure S3A-G**). Furthermore, we evaluated two additional mRNA expression datasets from TCGA (**Figure S4A**) and ICGC (**Figure S4B**) with the results revealing similar overexpression of these five hub genes in ccRCC. In addition, expression of these 5 hub genes was positively correlated with ccRCC stage (**Figure S4C**), grade (**Figure S4D**), T (thickness) and M (metastatic) classifications (**Figure S4E, 4F**). Thus, high expression of the 5 hub genes correlates with ccRCC progression.

Kaplan-Meier survival analysis was performed to evaluate the prognostic value of the five hub genes. High expression of all 5 genes was negatively correlated with overall survival (OS) and disease-free survival (DFS) (**Figure 1B, C, S5A-5H**). Univariable Cox regression analysis showed that all five hub genes were independent variables in predicting the clinical outcome of ccRCC patients (**Figure 1D**). Characteristics of the ccRCC patients from TCGA included in this study are shown in Supplementary Data File 1.

### NDC80 overexpression drives the malignant phenotype of ccRCC

Further mining of orthogonal gene expression datasets from the TCGA, TARGET and GTEx ^21,22^ databases revealed high expression of NDC80 in 30 out of 34 major human cancer entities (**Figure 2A**), an observation that is consistent with previous reports (^13–16^). To further illustrate the overexpression of NDC80 in ccRCC, we obtained from the TCGA and ICGC databases RNA expression data of ccRCCs paired with normal kidney tissue from the same patients. Comparative analysis of the paired samples revealed highly significant overexpression of NDC80 mRNA in ccRCC samples from TCGA (*P* = 1.4e-14) and ICGC (*P* = 1.2e-11) (**Figure 2B, C**). To determine NDC80 protein expression, we performed immunohistochemistry staining of 15 pairs of ccRCCs and matched normal tissues. Tumors showed strongly increased staining for NDC80 compared with normal kidney tissue (*P* < 0.0001) (**Figure 2D, E**). Likewise, immunoblotting showed that NDC80 protein levels were higher in the human ccRCC cell lines 786-O, 769-P and Caki-1 than in HK2 normal kidney cells (**Figure 2F, G**).

**Figure 2.**
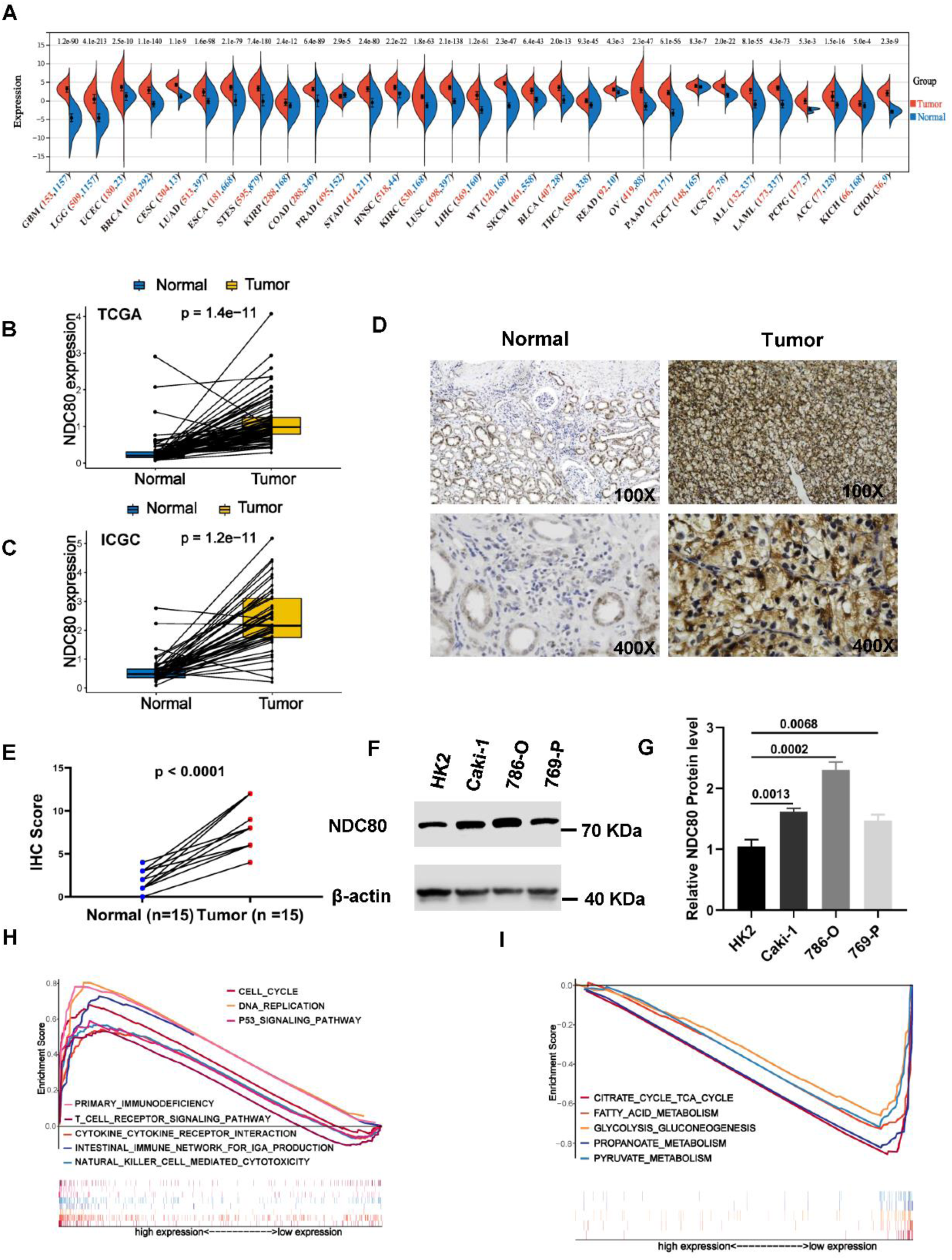
Expression of NDC80 in ccRCC. (**A)** NDC80 mRNA expression levels in 34 types of cancer based on TCGA, TARGET and GTEx datasets. (**B, C)** NDC80 mRNA expression level in 45 (TCGA) and 72 (ICGC) paired samples of ccRCC tissues and adjacent normal tissues. (**D)** Expression of NDC80 in paired ccRCC and normal kidney samples as determined by immunohistochemistry. A representative example is shown. (**E)** NDC80 immunostaining score in paired ccRCC and normal kidney samples (n = 15, averages +/− standard deviation). (**F, G)** Immunoblotting analysis of NDC80 protein expression in HK2, Caki-1, 786-O and 769-P cells. The protein expression of β-actin is shown as a reference (n = 6, averages +/− standard deviation). (**H, I)** KEGG pathways enriched in differential mRNA profiles of ccRCCs with high (left) or low (right) NDC80 expression as determined by GSEA.

To delineate the potential functions of NDC80 in ccRCC, we determined KEGG pathways differentiating NDC80-low and NDC80-high cancers by GSEA. Highly expressed gene sets in NDC80-high ccRCC included cell cycle, cytokine receptor interaction, DNA replication, p53 signaling pathway, intestinal immune network, natural killer cell-mediated cytotoxicity, primary immunodeficiency, and T cell receptor signaling pathway (**Figure 2H**). In contrast, TCA cycle, fatty acid metabolism, and glycolysis/gluconeogenesis pathways were downregulated in NDC80-low ccRCCs (**Figure 2I**).

To experimentally underpin these functional enrichments, NDC80 was knocked down in 786-O and Caki-1 ccRCC cells with two different siRNAs whereby siRNA1 was consistently less efficient than siRNA2 (**Figure 3A**). As shown by CCK8 assay, knockdown of NDC80 inhibited the proliferation of both cell lines (**Figure 3B**).

**Figure 3.**
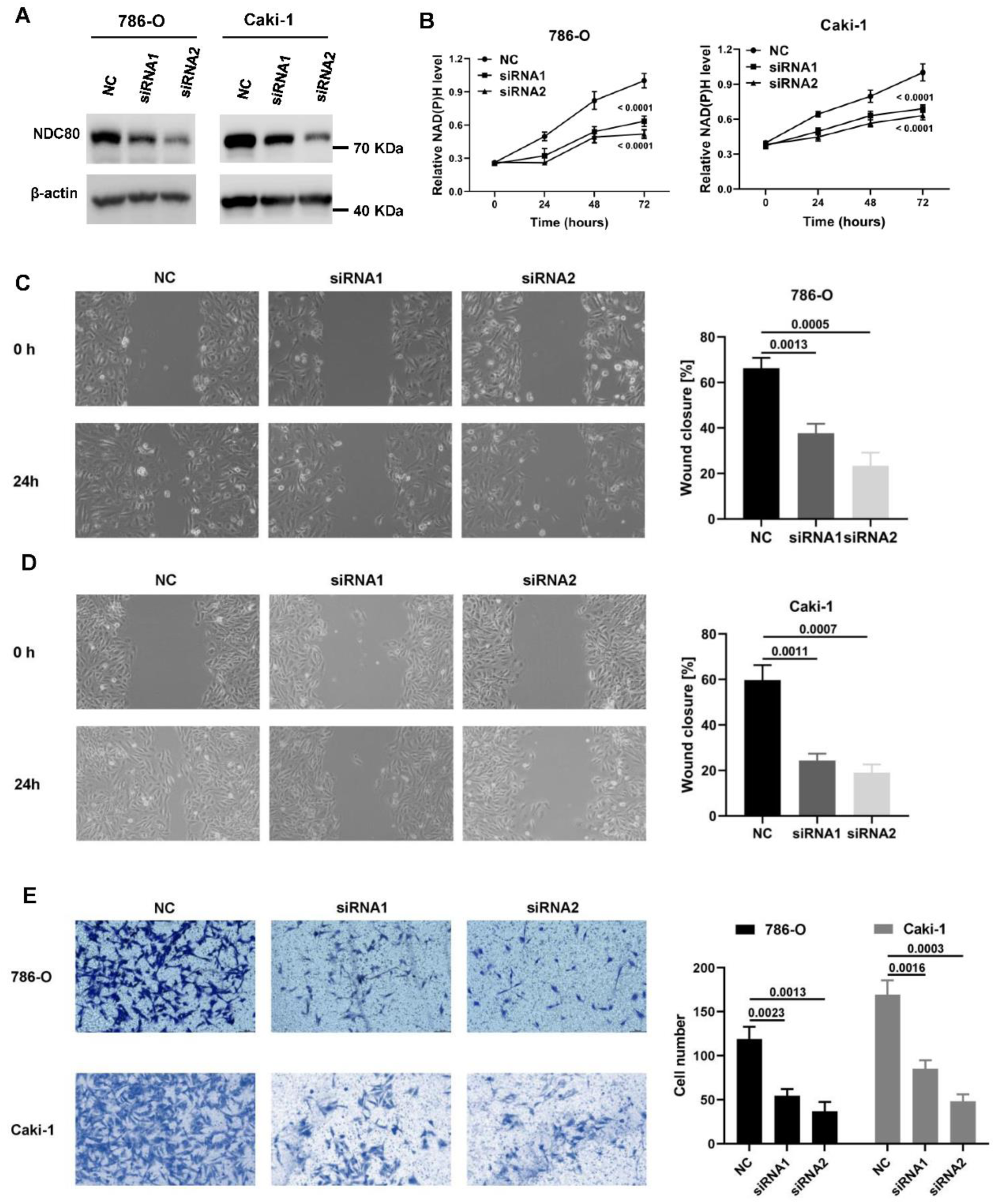
Effect of NDC80 on cell proliferation, migration, and invasion. (**A)** Total lysates prepared from 786-O and Caki-1 cells transfected with control or two different NDC80 siRNAs were assayed by immunoblotting with antibodies against NDC80. The signal obtained with β-actin antibodies is shown as a reference. (**B)** NAD(P)H levels as a surrogate measure of cell proliferation were measured in 786-O and Caki-1 cells transfected with control siRNA or with two different NDC80 siRNAs (n = 3, averages +/− standard deviation). (**C, D)** Wound healing assay was performed to measure migration of 786-O and Caki-1 cells after NDC80 knockdown (n = 3, averages +/− standard deviation). (**E)** Transwell assay was performed to measure invasion of 786-O and Caki-1 cells after knockdown of NDC80 (n = 3, averages +/− standard deviation).

NDC80 knockdown inhibited cell migration and invasion of 786-O and Caki-1 cells, two hallmarks of the malignant phenotype (**Figure 3C, D, E**). Flow cytometric measurement of DNA content revealed that NDC80 knockdown cells accumulated in S phase (**Figures 4A**), suggesting that NDC80 deficiency caused a cell cycle arrest in S phase, which is consistent with previous findings in liver cancer cells ^23^.

**Figure 4.**
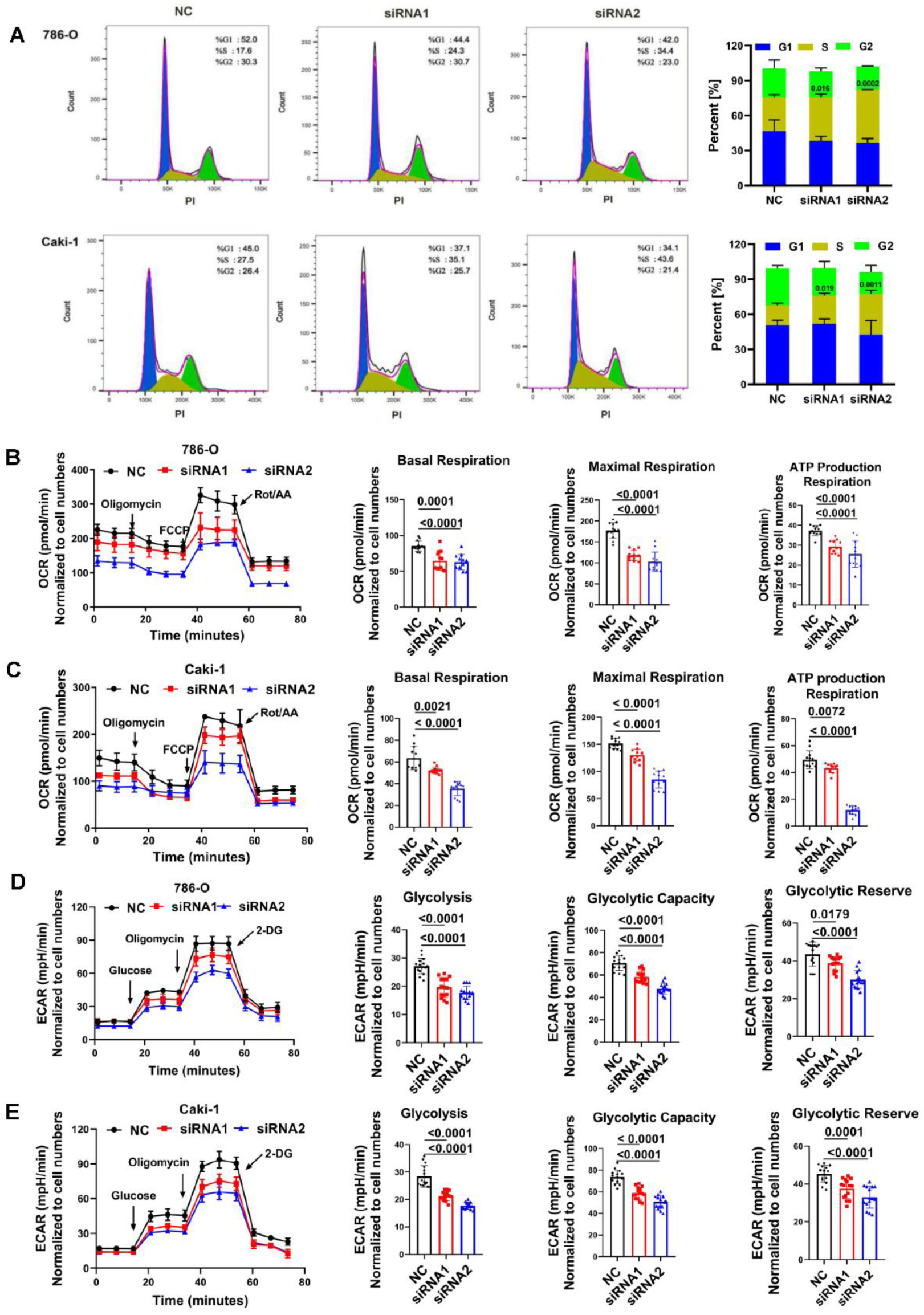
Effect NDC80 on cell cycle and metabolism. (**A)** 786-O and Caki-1 cells were transfected with NDC80 siRNA and control siRNA for 72 hours and cell cycle distribution of cells stained with propidium iodide was determined by flow cytrometry (n = 3, averages +/− standard deviation). (**B, C)** 786-O and Caki-1 cells were transfected with NDC80 siRNA and control siRNA for 48 hours and oxygen consumption rate (OCR) was measured in the Agilent Seahorse XFe96 metabolic flux analyzer (n = 18, averages +/− standard deviation). (**D, E)** Same as above but extracellular acidification rate (ECAR) was measured (n = 18, averages +/− standard deviation).

Furthermore, to test the prediction that NDC80 affects metabolism, in particular TCA cycle and glycolysis, we measured oxygen consumption rate (OCR) and extracellular acidification rate (ECAR) after NDC80 knockdown in 786-O and Caki-1 cells (**Figure 4B, C**). In both cell lines, basal and maximal respiration rates were reduced along with ATP production (*P* < 0.05). Likewise, glycolysis, including glycolytic capacity and glycolytic reserve, was significantly reduced after NDC80 knockdown in 786-O and Caki-1 cells (**Figure 4D, E**).

Finally, we sought to determine whether overexpression of NDC80 in NDC80-low HK2 cells would dominantly trigger events that depended on NDC80 in NDC80-high 786-O and Caki-1 cells. Remarkably, transient overexpression of NDC80 (Figure 5A) was sufficient to stimulate the proliferation, cell migration, invasion, mitochondrial respiration, and glycolysis of HK2 cells (Figure 5B-E). Thus, as predicted by network analysis and GSEA (**Figures 1A, 2H, I**), our data suggest that NDC80 has an unanticipated role in driving cell proliferation and central carbon metabolism of ccRCC cells.

**Figure 5.**
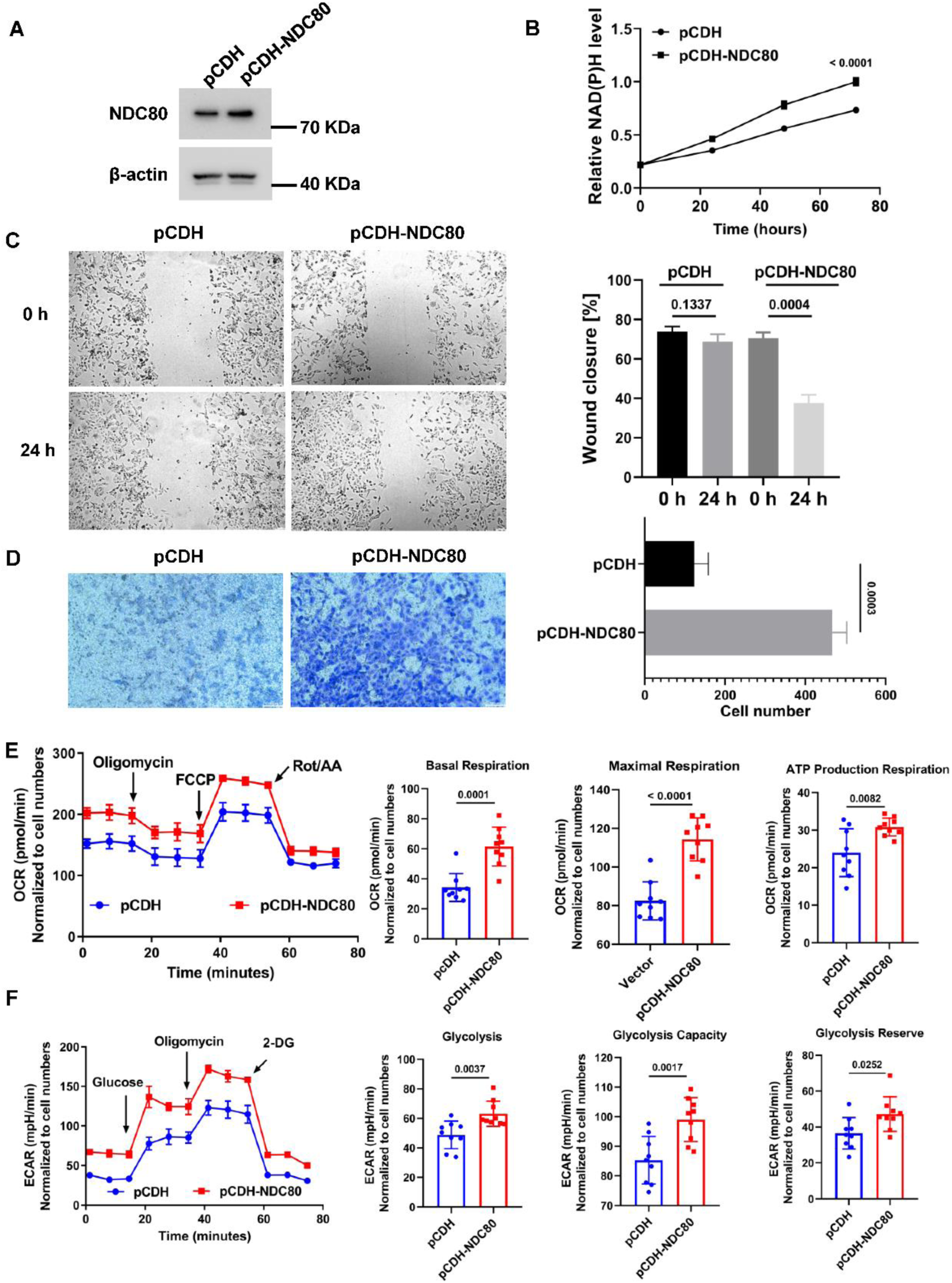
Effect of ectopic expression of NDC80 on HK2 cells. (**A)** Total lysates prepared from HK2 cells transiently transfected with a plasmid driving the expression of NDC80 for 48 hours were assayed by immunoblotting with antibodies against NDC80. The signal obtained with β-actin antibodies is shown as a reference. (**B)** NAD(P)H levels as a surrogate measure of cell proliferation were measured in HK2 cells overexpressing NDC80 (n = 4, averages +/− standard deviation). (**C)** Wound healing assay was performed to measure migration of NDC80 overexpressing HK2 cells (n = 3, averages +/− standard deviation). (**D)** Transwell assay was performed to measure invasion of NDC80 overexpressing HK2 cells (n = 3, averages +/− standard deviation). (**E)** HK2 cells were transiently transfected with a plasmid driving the expression of NDC80 for 48 hours and oxygen consumption rate (OCR) was measured in the Agilent Seahorse XFe96 metabolic flux analyzer (n = 9, averages +/− standard deviation). (**F)** Same as above but extracellular acidification rate (ECAR) was measured (n = 9, averages +/− standard deviation).

### High NDC80 predicts poor response to immunotherapy

The GSEA results suggested that NDC80 expression affects immune cell function. Using the ESTIMATE algorithm, we found that the immune score (*P* = 1e^−10^), stromal score (*P* = 2.9e^−6^), and ESTIMATE score (*P* = 9.3e^−11^) were higher in the NDC80-high versus the NDC80-low group (**Figure 6A, Figure S6A, S6B**) thus validating the GSEA findings. Further analysis with the TIMER algorithm ^24^, a comprehensive resource for systematic analysis of immune infiltrates from bulk RNA-seq data, indicated that NDC80 expression was positively correlated with the abundance of B cells (R = 0.282, *P* = 3.95e^−11^), CD4+ T cells (R = 0.327, *P* = 1.02e^−14^), CD8+ T cells (R = 0.435, *P* = 7.3e^−26^), neutrophils (R = 0.392, *P* = 5.53e^−21^), macrophages (R = 0.244, *P* = 1.22e^−8^), and dendritic cells (R = 0.282, *P* = 9.67e^−28^) (**Figure 6B**). The immune associations were confirmed with a related algorithm to impute gene expression profiles, CIBERSORT, which indicated that high NDC80 expression is strongly correlated with 12 classes of tumor immune cells, including several types of CD4+ and CD8+ T cells, NK cells, monocytes, macrophages, dendritic cells, and mast cells (**Figure 6C**). The imputations were further confirmed by the analysis of single cell RNA-seq data (GSE152938 and GSE171306), which showed a high level of immune infiltration of ccRCCs (**Figure 6D**). In addition, this analysis revealed that NDC80 is highly expressed in T cells, especially proliferative CD8+ T cells which were described as dysfunctional ^25,26^ (**Figure 6E**). NDC80-high proliferative CD8+ T cells were also consistently positive for well-established markers of exhaustion, including TIGIT, PDCD1, TOX, RGS1, and LAG3 (**Figure 6E**).

**Figure 6.**
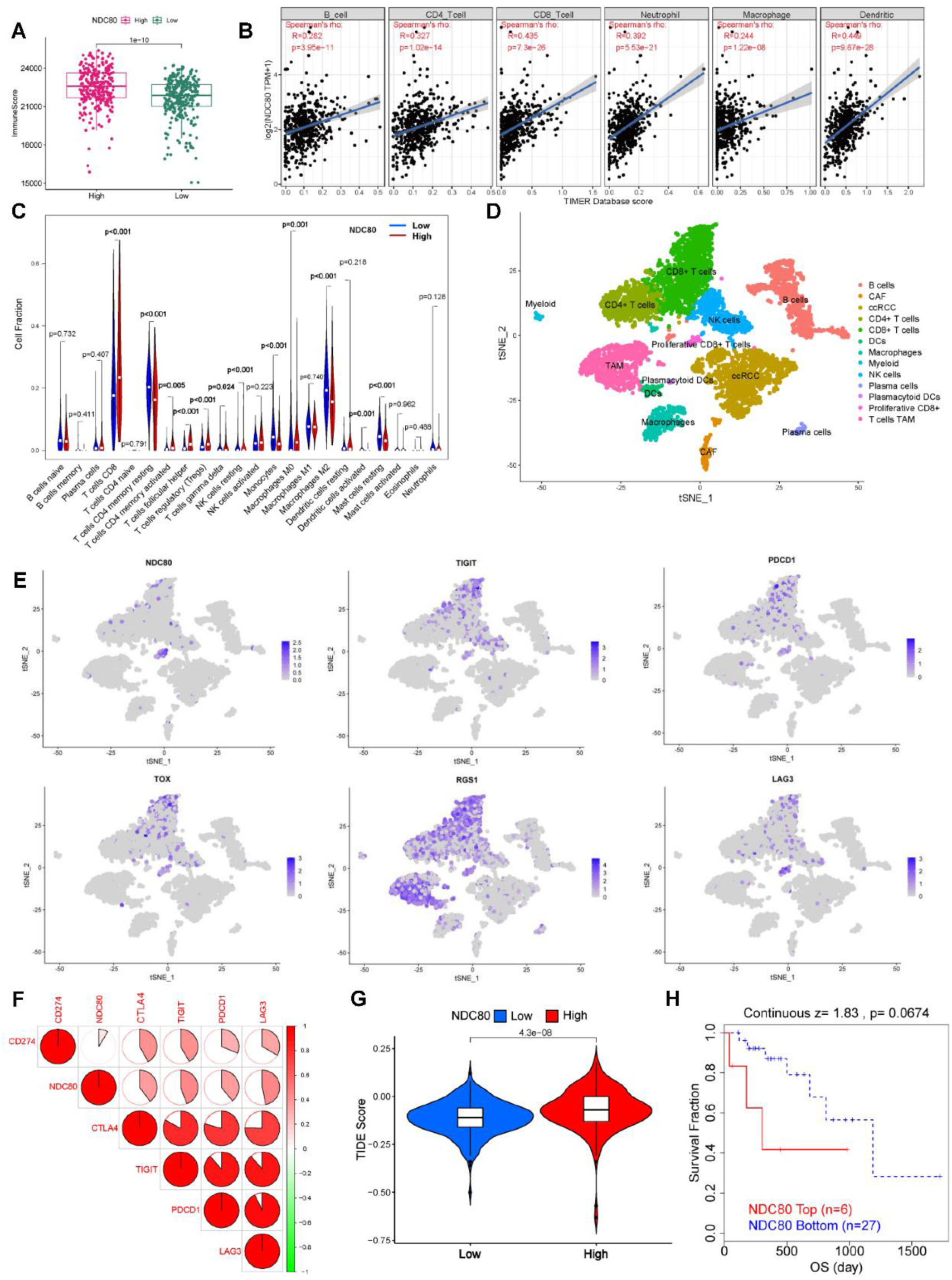
Correlation of NDC80 expression with immune infiltration and response to immunotherapy in ccRCC. (**A)** ESTIMATE algorithm was used to calculate immune scores in NDC80-low and NDC80-high ccRCC mRNA expression datasets from TCGA. (**B)** TIMER algorithm was performed to calculate the correlation between NDC80 expression and abundance of the indicated immune cells. (**C)** CIBERSOFT algorithm was used to correlate NDC80 expression with the fraction of 22 immune cells. (**D)** Annotation of virous cell types in four ccRCC single cell mRNA sequencing datasets obtained from GEO. (**E)** NDC80 expression in different cell types was determined with the feature plot function using R package “Seurat”. (**F)** Spearman correlation between expression of NDC80 and immune checkpoint factors CD274 (PDL1), PDCD1 (PD1), TIGIT, CTLA4 and LAG3 in the KIRC dataset from TCGA. (**G)** Tumor Immune Dysfunction and Exclusion (TIDE) algorithm was performed to assess the TIDE score for NDC80-high and NDC80-low ccRCC samples. (**H)** Kaplan–Meier plots of overall survival (OS) of ccRCC patients treated with PD1 antibody from TIDE.

Further consistent with NDC80 indicating T cell dysfunction was the observation that NDC80 is positively correlated with the expression of immune checkpoint regulators including PD1, PD-L1, CTLA4, LAG3 and TIGIT (**Figure 6F**). Indeed, analysis with the Tumor Immune Dysfunction and Exclusion (TIDE) algorithm ^27^ showed that NDC80-high ccRCCs have a higher TIDE score than the NDC80-low group, indicating that NDC80-high patients have a worse response to immunotherapy (**Figure 6G**). The OS data of ccRCC patients treated with PD1 antibody from TIDE confirmed that patients with NDC80-high ccRCCs had poorer survival in response to PD-1 antibody treatment than patients with NDC80-low tumors (**Figure 6H**). Thus, despite correlating with a high immune score, aggressive NDC80-high ccRCCs respond poorly to immunotherapy thus creating a strong need for alternative therapeutic avenues.

### NDC80 predicts sensitivity of ccRCCs to mitotic kinase inhibition

In attempts to identify drugs that might be able to target NDC80-high ccRCCs, we mined the Genomics of Drug Sensitivity in Cancer (GDSC2) database ^28^ for correlations between drug response (IC_50_) and NDC80 expression in ccRCC cell lines. 124 compounds showed significant correlations (*p* < 0.05) with NDC80 expression. IC_50_ values of the majority of compounds (111) were positively correlated with NDC80 levels, suggesting that NDC80-high cancer cell lines are generally less responsive to drug treatment than NDC80-low cell lines (**Supplementary Data File 2**). However, 13 compounds showed a negative correlation, indicating preferential sensitivity of NDC80-high cell lines. The 3 most highly correlated compounds included tozasertib (R = −0.37, p = 0.0105) and ZM447439 (R = −0.29, p = 0.0458), two inhibitors of aurora kinases (AURKA/B/C) known to be involved in the spindle assembly checkpoint and chromosome segregation (**Figure 7A, B**). An equally high correlation was obtained for BI-2536 (R = −0.34, p = 0.0271), an inhibitor of another mitotic regulator, polo kinase 1 (PLK1) (**Figure 7C**).

**Figure 7.**
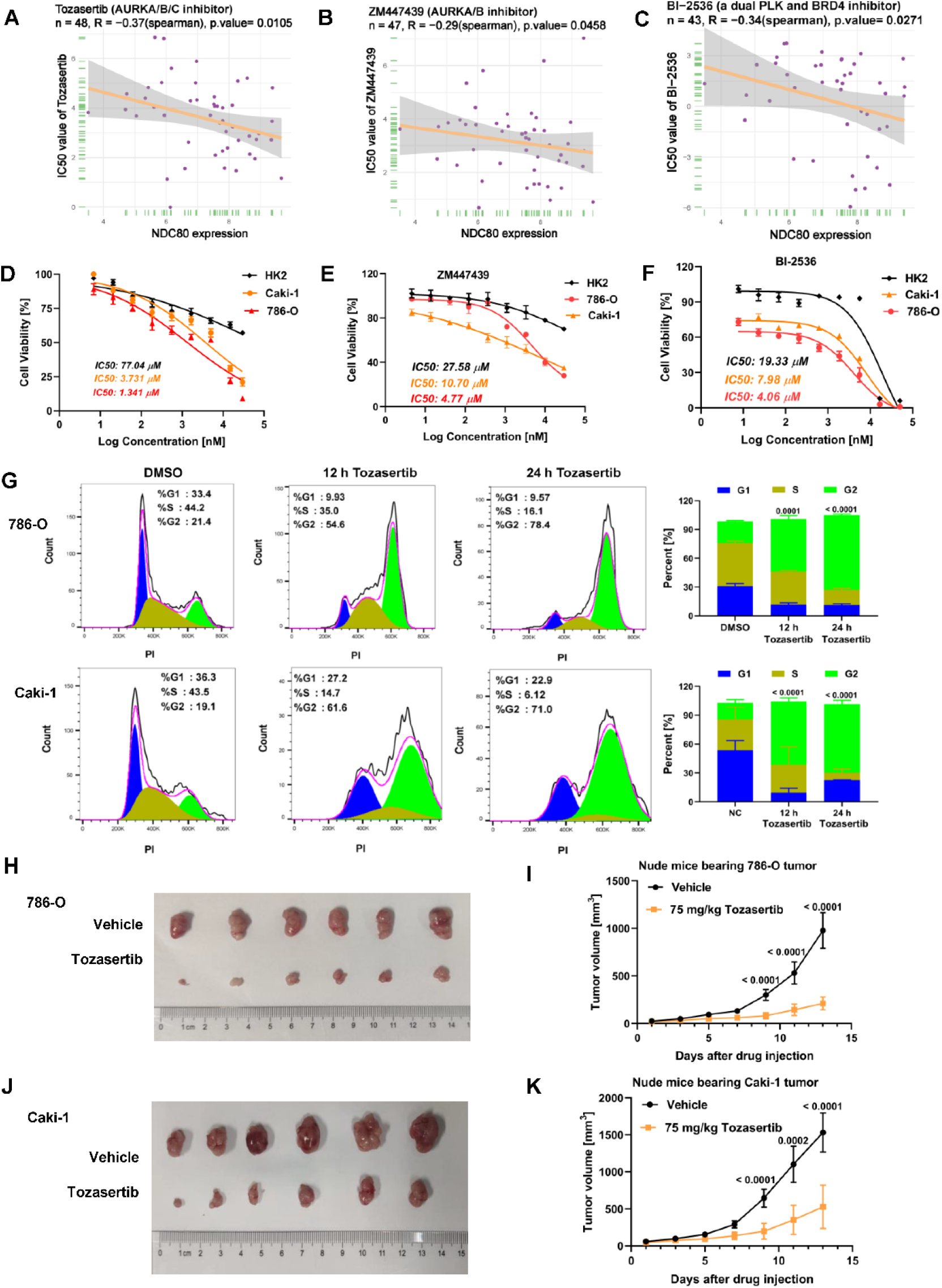
Correlation of NDC80 expression with sensitivity of ccRCCs to mitotic kinase inhibition *in vitro* and *in vivo*. (**A-C)** Correlation between the IC_50_ of tozasertib (AURKA/B/C inhibitor), ZM447439 (AURKA/B inhibitor), and BI-2536 (PLK and BRD4 inhibitor) and NDC80 expression in cancer cell lines. Data from the Genomics of Drug Sensitivity in Cancer (GDSC). (**D-F)** Effect of the indicated inhibitors on metabolic activity as a surrogate of cell proliferation in HK2, 786-O and Caki-1 cells (n = 6, averages +/− standard deviation). (**G)** Effect of DMSO or tozasertib treatment for 12- or 24-hours on cell cycle distribution in 786-O and Caki-1 cells. The cell lines were treated with a concentration of tozasertib equivalent to their respective IC_50_ (n = 3, averages +/− standard deviation). (**H-K)** 786-O and Caki-1 cells were grown as tumors in nude BALB/c mice. Mice were treated with 75 mg/kg tozasertib or vehicle by i.p. every 48 hours, and tumor growth was followed for 2 weeks (n = 6, averages +/− standard deviation).

To assess the effect of these inhibitors on ccRCC cells, we compared their dose responses in 786-O and Caki-1 cells relative to HK2 normal kidney cells. ccRCC cell lines were between 57- and 20-fold more sensitive to tozasertib, between 5.8- and 2.6-fold more sensitive to ZM447439 and between 4.8- and 2.4-fold more sensitive to BI-2536 than HK2 cells (**Figure 7D, E, F**), suggesting a substantial therapeutic index. Consistent with their targeting of mitotic kinases, all 3 compounds induced cell cycle arrest in G2/M phase (**Figure 7G, Figure S7A, B**).

Finally, we assessed the anti-tumor activity of tozasertib in mouse xenografts of 786-O and Caki-1 tumors. Injection of 75 mg/kg tozasertib i.p. for two weeks every 48 hours significantly reduced the growth of 786-O and Caki-1 tumors over time (**Figure 7H, I**). Average tumor sizes at the end of the treatment period were also substantially reduced (**Figure S7C, S7D**), whereas body weights did not change (**Figure S7E, S7F**). Thus, inhibitors of mitotic kinases such as tozasertib show promise for safely targeting hard-to-treat NDC80-high ccRCCs. Tozasertib treatment of 786-O and Caki-1 cells for up to 24 hours did not affect the levels of NDC80 (**Figure S7G**).

## DISCUSSION

In this study, we performed unbiased bioinformatic analyses to identify NDC80 as an important prognostic marker in ccRCC. Not only does high expression of NDC80 predict poor patient outcome, but it specifically pinpoints cancers with a poor response to immunotherapy. On the flipside, high NDC80 expression predicts a favorable response to inhibition of mitotic regulatory kinases, AURKA/B and PLK1/2.

NDC80’s predictive power may derive from its function in driving the malignant phenotype. As such we found that NDC80 is overexpressed in the majority of human cancers, a finding that is consistent with numerous previous studies on individual cancer entities (e.g. ^15,16,23^). Overexpression of NDC80 in transgenic mice causes hyperactivation of the spindle assembly checkpoint, accumulation of cells in mitosis, chromosome instability, and tumors marked by aneuploidy ^13^. Similarly, depletion of NDC80 leads to accumulation of various cancer cells in G2/M ^15,16^. In ccRCC cell lines, however, we found that depletion of NDC80 causes accumulation of cells in S phase rather than G2/M (**Figure 3I**), indicating that NDC80’s tumor promoting activity may not be restricted to controlling mitotic fidelity but directly drive cell proliferation. A similar S phase advancing activity of NDC80 was previously described in hepatocellular carcinoma cells ^23^. It is noteworthy in this regard that several S phases promoting factors, including cyclins A2 (CCNA2), D1 (CCND1), and E2 (CCNE2) as well as CDC6 are components of the NDC80 PPI sub-network (**Figure S3A**). It is thus plausible that NDC80 impacts cell proliferation through multiple cell cycle regulatory events. For example, NDC80 might be involved in the licensing of replication origins, although this remains to be tested.

Our bioinformatic analyses pointed to several additional pathways for how NDC80 overexpression might drive tumor progression and treatment resistance. GSEA indicated a positive correlation between NDC80 expression and metabolic pathways, including glycolysis and mitochondrial respiration which we confirmed experimentally (**Figure 4B, C, D, E, 5E, F**). Considering the frequent inactivation of VHL and accumulation of HIF1 transcription factors in ccRCC, a positive association of NDC80 with glycolysis seems unsurprising, whereas the positive association with mitochondrial respiration was unexpected. However, high mitochondrial respiratory activity (“OXPHOS”) has been linked with NDC80 driven resistance to cisplatin ^29^ while knockdown of NDC80 increased sensitivity of cancer cells to paclitaxel ^14^. Our analysis of GDSC data has also indicated widespread resistance of NDC80-high cells to a large variety of anti-cancer agents (**Supplementary Data File 2**). In addition, a high rate of oxygen consumption by the tumor cells is known to augment tumor hypoxia and could thus further boost HIF1α upregulation while at the same time causing T cell exhaustion ^30,31^. The latter conjecture is consistent with our analysis of the immune phenotype of NDC80-high ccRCCs which indicated substantial infiltration by T cells with an exhausted phenotype (**Figure 6E**). Through these complex connections, NDC80 driven mitochondrial respiration, albeit mechanistically still unexplained, might contribute to compromised tumor immunity thus leading to poor response to immunotherapy.

While high NDC80 levels coincided with broad therapy resistance, it correlated with increased sensitivity to inhibitors of AURK as well as PLK1. Inhibition of both kinases, which are intertwined in a multifaceted network controlling many aspects of mitosis, including centrosome maturation and separation, mitotic entry, spindle assembly, kinetochore–microtubule attachment, the spindle assembly checkpoint, DNA damage checkpoint activation, and cytokinesis ^32,33^, led to inhibition of ccRCC proliferation through induction of a G2/M phase cell cycle delay (**Figure 6G, S7A, S7B**). Notably, PLK1 is known to activate AURKB which in turn phosphorylates NDC80, thereby regulating its interaction with microtubules. It is thus possible that this direct enzyme-substrate cascade instigates the preferential sensitivity of NDC80-high ccRCCs.

Since knockdown of NDC80 induces an S phase arrest, whereas mitotic kinase inhibitors instigate a G2/M arrest, the kinase inhibitors do not appear to impact the viability of ccRCC cells simply by inhibiting NDC80 function. Mitotic kinase inhibition likely affects the activity of many proteins in addition to NDC80, whereas siRNA-mediated knockdown is relatively specific in disabling NDC80. Our data suggest that NDC80 has a function in S phase that is not controlled by mitotic kinases but which stimulates the proliferation of ccRCC cells. At the same time, high NDC80 levels may promote high mitotic activity that confers sensitivity to AURK and PLK inhibition.

Two of the compounds active against ccRCCs, AURK inhibitor tozasertib and PLK1 inhibitor BI-2536 and its second-generation analog BI-6727 (volasertib) underwent numerous phase I and II clinical trials against non-solid and solid tumors but not ccRCC ^34^. The compounds were marked by limited efficacy as monotherapies in unselected patient cohorts with further clinical development of BI-6727 awaiting the identification of biomarkers predicting clinical response as those investigated to date (PLK1, Ki-67, AURKA/B, and pHH3) were not effective ^35^. Importantly, our studies suggest NDC80 as the first function-based biomarker of sensitivity to mitotic kinase inhibitors in ccRCC and possibly other cancers. If further validated, inclusion of NDC80 into clinical trial design may clear the path to continued development of mitotic kinase inhibitors as treatments for ccRCC and other malignancies.

## LIMITATIONS OF THE STUDY

Although we have experimentally validated some of the key predictions derived from the in-silico data mining – e.g., the role of NDC80 in proliferation, cell cycle progression, and cell migration as well as the sensitivity of NDC80-high tumors to mitotic kinase inhibitors – these studies rely on a rather limited set of cellular models. In addition, other aspects of our study will require additional experimental validation. This primarily pertains to the potential immune modulatory effects of NDC80 which appear to be due to complex microenvironmental control that we were unable to fully resolve in the present study. Finally, the potential applicability - for example with respect to sensitivity and specificity - of NDC80 as a biomarker predicting the response of ccRCC patients to mitotic kinase inhibitors will need to be resolved in future clinical studies.

## ACKNOWLEDGMENTS

The support from the Equipment Platform of the State Key Lab of Cellular Stress Biology at Xiamen University is gratefully acknowledged. This work was supported by National Natural Science Foundation of China (81773771, D.A.W), Fujian Province Science and Technology projects (2018J01053, 2021Y0002, Y.C.), and Fundamental Research Funds for the Central Universities (20720220053, Y.C.). Additional support was from hospital funds (G. L., X. K.).

## AUTHOR CONTRIBUTIONS

Conceptualization, C.H., D.A.W, Y.C.; Methodology, C.H., W.L., K.Z., G.T.; Formal Analysis, C.H., W.L., K.Z., Y.Z.; Investigation, C.H., W.L., K.Z., G.T.; Writing –Original Draft, C.H., D.A.W., Y.C.; Writing – Review & Editing, C.H., W.L., K.Z., G.T., D.A.W., Y.C., G.L., X.K.; Visualization, C.H., W.L., K.Z., G.T.; Supervision, D.A.W., Y.C.; Funding Acquisition, D.A.W., Y.C., G.L., X.K..

## STAR★METHODS

### KEY RESOURCES TABLE

| REAGENT or RESOURCE | SOURCE | IDENTIFIER |
| --- | --- | --- |
| **Deposited data** |  |  |
| TCGA-KIRC | The Cancer Genome Atlas (TCGA) <sup>36</sup> | https://portal.gdc.cancer.gov/ |
| ICGC | ICGC Data Portal <sup>37</sup> | https://dcc.icgc.org/ |
| GSE36895 | Peña et al., <sup>37</sup> | https://www.ncbi.nlm.nih.gov/geo/ |
| GSE53000 | Gerlinger et al., <sup>38</sup> | https://www.ncbi.nlm.nih.gov/geo/ |
| GSE53757 | Von Roemeling et al., <sup>39</sup> | https://www.ncbi.nlm.nih.gov/geo/ |
| GSE66272 | Wotschofsky et al., <sup>40</sup> | https://www.ncbi.nlm.nih.gov/geo/ |
| GSE152938 | Su et al., <sup>41</sup> | https://www.ncbi.nlm.nih.gov/geo/ |
| GSE171306 | Huang et al., <sup>42</sup> | https://www.ncbi.nlm.nih.gov/geo/ |
| Pan-cancer expression | UCSC Xena <sup>22</sup> | https://xenabrowser.net/hub/ |
| Protein expression of hub genes | UALCAN <sup>20</sup> | http://ualcan.path.uab.edu/analysis-prot.html |
| **Software and Algorithms** |  |  |
| R (v 4.1.3) |  |  |
| ESTIMATE | Yoshihara et al., <sup>43</sup> | https://bioinformatics.mdanderson.org/estimate/ |
| TIMER | Li et al., <sup>24</sup> | https://cistrome.shinyapps.io/timer/ |
| CIBERSOFT | Newman et al., <sup>44</sup> | https://cibersortx.stanford.edu/ |
| TIDE | Fu et al., <sup>27</sup> | http://tide.dfci.harvard.edu/ |

| Reagent |  |  |
| --- | --- | --- |
| CCK-8 | Absin | Abs50003 |
| PI | Sigma-Aldrich | 81845 |
| RNase A | Solarbio | R1030 |
| Matrigel | BD Biosciences | 356234 |
| Crystal violet | Sigma-Aldrich | C0775 |
| Tozasertib | MedChemExpress | HY-10161 |
| BI-2536 | MedChemExpress | HY-50698 |
| ZM447439 | MedChemExpress | HY-10128 |
| Oligomycin | MedChemExpress | HY-N6782 |
| FCCP | MedChemExpress | HY-100410 |
| Rotenone | Sigma-Aldrich | 45656 |
| Antimycin A | Sigma-Aldrich | A8674 |
| 2-DG | Sigma-Aldrich | D8375 |
| D-glucose | Sigma-Aldrich | G8270 |
| Protease Inhibitor Cocktail | ThermoFisher | 78438 |
| Anti-NDC80 | Proteintech | 18932-1-AP |
| Anti-β-actin | ABclonal | AC006 |
| HRP Goat Anti-Rabbit IgG (H+L) | ABclonal | AS014 |

### RESOURCE AVAILABILITY

#### Lead contact

Further information and requests for resources should be directed to and will be fulfilled by the lead contact, Yabin Cheng (

#### Materials availability

This study did not generate new unique reagents or material.

### EXPERIMENTAL MODEL AND SUBJECT DETAILS

#### Data collection

The data of the ccRCC cohorts were downloaded from The Cancer Genome Atlas (TCGA) ^36^ and Gene Expression Omnibus (GEO) databases. Gene expression data were acquired from TCGA-KIRC and GSE53757, GSE36895, GSE66272, GSE53000 ^38–40,45^, while the clinicopathological and prognostic information was obtained from TCGA KIRC cohort. Cases without detailed clinical information or lacking gene expression data were omitted. Protein expression data were obtained from UALCAN^20^. RNA-seq data on 72 and 45 paired tumors and controls were obtained from the TCGA and ICGC databases ^37^, respectively. The single cell RNA-seq data of four ccRCC samples was collected from GEO (GSE152938 and GSE171306) ^41,46^. For NDC80 expression in pan-cancer analysis, we downloaded the normalized pan-cancer dataset from the UCSC database ^22^: TCGA, TARGET, Genotype-Tissue Expression (GTEx) ^21^(PANCAN, N = 19131, G = 60499). Cancer species with less than 3 samples were eliminated and then NDC80 gene expression data was extracted in each sample. Each expression value was transformed using log2 (x + 0.001).

#### Differential gene expression (DEGs) and pathway enrichment analysis

The R package “limma” was run to identify differentially expressed genes (Log2 Fold Change > 1, *P* value < 0.05) in the GSE53757, GSE36895, GSE66272 and GSE53000 cohorts. Lists of 338 upregulated and 405 downregulated genes common to all four cohorts were obtained using the R package “Venn”. Gene oncology (GO) Biological Process and Kyoto Encyclopedia of Genes and Genomes (KEGG) enrichment analysis was performed by using the R package ‘clusterProfiler’.

#### Identification of hub genes and protein-protein interactions

Protein-protein interactions (PPI) corresponding to the proteins encoded by the 743 DEGs were loaded into the STRING database (https://string-db.org/) ^47^. After adjusting the minimum required interaction score to 0.7, a .tsv file was exported from STRING. The PPI network was loaded into Cytoscape (11.0.6 version). The top ten hub genes/proteins were obtained based on their ranking by the Maximal Clique Centrality (MCC) method using the Cytohubba app.

### METHOD DETAILS

#### mRNA and protein expression levels of hub genes based on TCGA and UALCAN

To explore the protein expression level of the 10 hub genes, liquid chromatography tandem mass spectrometry (LC-MS/MS) data of ccRCC samples was obtained from the Clinical Proteomic Tumor Analysis Consortium (CPTAC) in UALCAN (http://ualcan.path.uab.edu/analysis-prot.html). Only 7 of the 10 hub proteins (ASPM, CDK1, NCAPG, NDC80, PBK, KIF11 and CCNA2) were detected in ccRCC samples by LC-MS/MS. The mRNA expression of CDK1, NCAPG, ASPM, NDC80 and PBK were visualized using R packages “ggplot2”.

#### Survival analysis

Survival analysis was performed by Kaplan-Meier plots of overall survival (OS) and disease-free survival (DFS) data obtained from the TCGA database. Univariable Cox regression analysis was performed to evaluate the prognostic value of NDC80. The cut-off value for OS and DFS analysis was determined by the median value of NDC80 expression.

#### Gene set enrichment analysis

The gene set enrichment analysis program (GSEA_4.0.3 version) was downloaded from the Gene Set Enrichment Analysis website (http://software.broadinstitute.org/gsea/index.jsp) ^47^. The input files of NDC80-high and NDC80-low samples were prepared in R. Based on the default weighted enrichment statistical method, the procedure was iterated 1000 times for each analysis. The signaling pathways with FDR < 0.05 were chosen as significantly regulated pathways.

#### ESTIMATE, TIMER, CIBERSORT and TIDE analysis

ESTIMATE ^43^ and TIMER ^24^ are comprehensive resources for systematic analysis of immune infiltrates across diverse cancer types (https://cistrome.shinyapps.io/timer/). The algorithms apply statistical deconvolution methods to infer the abundance of tumor-infiltrating immune cells from gene expression profiles. We analyzed the correlation of NDC80 expression with immune infiltrates including B cells, CD8+ T cells, CD4+ T cells, neutrophils, macrophages, and dendritic cells via predefined gene modules. For more detailed analysis, the correlation of 22 types of immune cells with NDC80 expression was assessed with the CIBERSORT algorithm ^44^. The Tumor Immune Dysfunction and Exclusion (TIDE) ^27^ tool was used to calculate the TIDE score and collect the OS of ccRCC patients treated with immune checkpoint blockade therapy.

#### Single cell sequencing analysis

Single-cell RNA-seq data were from GSE152938 and GSE171306. Two ccRCC samples from GSE131685 and two ccRCC samples from GSE171306 were acquired from GEO. The four samples were merged for further analysis using R package “Seurat” ^48^. Single-cell RNA-seq data processing was performed as described ^49^. Cell clusters were annotated manually based on previous cell identity markers ^50^. The “FeaturePlot” function was used to show the expression of NDC80 and markers of T cell exhaustion in different cell clusters.

#### GDSC2 database analysis

The Genomics of Drug Sensitivity in Cancer database (GDSC) ^28^ was used to correlate drug dose responses (IC_50_) of 198 compounds with NDC80 expression in 809 cell lines. Correlations were analyzed by Spearman’s Rank Correlation Coefficient. 124 compounds showed significant correlations (*p* < 0.05) with NDC80 expression. 13 compounds were positively correlate with NDC80 expression, while 111 compounds showed a negative correlation.

#### Cell lines

Clear cell renal cell carcinoma cell lines (786-O, 769-P, Caki-1) and human renal proximal tubule epithelial cells (HK2) were obtained from FDCC (Shanghai, China). 786-O and 769-P cells were cultured in RPMI-1640 medium (Procell), Caki-1 was cultured in Myco-5A medium (Procell), while HK2 cell was cultured in MEM medium (Procell). All media was supplemented with 10% fetal bovine serum and 1% antibiotics. Cells were maintained in 5% CO_2_ at 37 °C.

#### Immunohistochemistry assay

15 clear cell renal cell carcinomas (HKid-CRC030PG-01) were obtained from Shanghai OUTDO Biotech. The immunohistochemistry staining procedure was done according to the Immunohistochemistry Protocol Paraffin for SignalStain® Boost Detection (Cell Signaling Technology). NDC80 antibody (Proteintech, 18932-1-AP) at a dilution of 1:200 was used for immunohistochemistry. The samples were scored into 5 bins based on the fraction of cells stained (positivity score): <10% = 0, 10 - 25% = 1, 25 - 50% =2, 50 - 75% = 3, 75 - 100% =4. The intensity of staining was scored into 4 four grades (intensity scores): no staining = 0, weak staining = 1, moderate staining = 2, strong staining = 3. NDC80 staining positivity was determined by the following formula: Overall score = positivity score × intensity score. An overall score ≤4 was defined as negative and a score > 4 was defined as positive.

#### DNA and RNA transfection

NDC80-targeting siRNA was obtained from Gene Pharma. The sequences of siRNA1, siRNA2, and unspecific control siRNA were 5’-GCAGCCUUAGUUUGGCUAATT-3’, 5’-GGAGGAUACUUUAGAACAATT-3’ and 5’-UUCUCCGAACGUGUCACGUTT-3’, respectively. Cells at ∼40% confluence were transfected in six well plates using Lipofectamine 2000 (Invitrogen, USA) according to the manufacturer’s instructions. After 48 hours, cells were collected to perform immunoblot analysis to document knockdown efficiency.

For ectopic expression, human NDC80 cDNA was cloned into the pCDH-CMV-MCS-EF1-copGFP-T2A-Puro plasmid resulting in pCDH-NDC80. Using Lipofectamine 2000, 293T cells were co-transfected with psPAX2 and pMD2.G to package pCDH-NDC80 for 48 h, and viral particles were filtered and frozen at −80°C. HK2 cells at a density of ∼50% to 60% were infected with viral particles and 4 μg/ml polybrene for 48 h after which cells were collected for further experiments.

#### Cell proliferation and cell cytotoxicity (CCK8) assay

After transfection with siRNAs or upon viral transduction of pCDH-NDC80, 2,000 cells per well were seeded into a 96 well plate in 100 μL medium. Cell Counting Kit-8 (Absin) was used to determine cellular metabolic activity (i.e. NAD(P)H levels) as a surrogate measure of cell proliferation. Cells were incubated with 10% CCK8 reagent diluted with fresh medium after 0, 24, 48 and 72 hours. After incubation for 1 hour, absorbance was measured at 450 nm using a microplate reader. Values were normalized to cells transfected with control siRNA and are represented graphically as mean ± standard deviation (STDEV) from three independent samples.

For compound cytotoxicity testing, 786-O (10,000) cells and Caki-1 (10,000) cells were seeded into 96 well plates. After 12 hours, 50 μM, 16.67 μM, 5.56 μM, 1.85 μM, 0.62 μM, 0.21 μM, 0.07 μM, 0.02 μM and 0 μM tozasertib (MCE), BI-2536 (MCE) or ZM447439 (MCE) were added into cells, respectively. After incubated for 48 hours, 10% CCK8 in fresh medium were added into cells and absorbance was measured as above.

#### Immunoblotting

Cells were lysed in RIPA buffer containing protease inhibitors on ice for 30 min, then centrifuged for 10 min at 10,000 g. Supernatants were taken for protein measurement with the BCA protein assay (Tiangen). Samples were boiled with SDS sample buffer for 10 min. Equal amounts of protein were separated by 10% SDS-PAGE and transferred to PVDF membranes (Merck Millipore). The membranes were incubated in 5% powdered nonfat milk in TBST for 1 hour, and incubation with primary antibodies was done overnight at 4 °C. Membranes were washed in TBST three times, followed by incubation with secondary antibodies (1:10,000; Abcam) for 1 hour at room temperature. After washing (TBST, 3x) immunoreactive products were detected by chemiluminescence and visualized and quantified with the ChemiDoc imaging system (Bio-Rad).

#### Wound healing and transwell assays

Cell migration was detected by wound healing assay. Cells were cultured in 6 well plates to 95% confluence. A 20 μL pipette tip was used to generate a scratch through each well. Wound closure was followed by microscopy (ZEISS). Scratch areas were measured with Image J software.

For invasion assays, 5 x 10^4^ transfected 786-O, Caki-1, or HK2 cells maintained in serum-free medium were seeded in the upper well of culture chambers equipped with 8 μm membranes (Corning Incorporated). The lower chamber was filled with 600 μL RPMI 1640 medium or Myco5A medium (Procell) supplemented with 10% FBS (Gibco) and 1% antibiotics (penicillin-streptomycin). After incubated at 37 °C, 5% CO_2_ for 24 hours, the upper chamber was washed with PBS 3 times, and cells that migrated to the lower surface of the chamber were fixed in 95% ethanol for 10 min, stained with 0.1% crystal violet solution for 10 min, washed with PBS 3x, air dried, photographed, and counted with ImageJ software.

#### Cell cycle analysis

After transfection with control and NDC80 siRNAs for 72 hours, cells were washed in PBS and fixed in 1 ml pre-cooled 70% ethanol for 4 hours in at −20 °C. Cells were pelleted at 200 x g for 5 min and suspended in 0.5 ml PBS containing 40 μg/ml propidium iodide solution and 100 μg/ml RNase A. Cells were incubated for 30 minutes at 37 °C and analyzed by flow cytometry.

#### Measurement of oxygen consumption rate (OCR) and extracellular acidification rate (ECAR)

Oxygen consumption rate (OCR) was measured utilizing the Agilent Seahorse XFe96 (Seahorse Bioscience, Billerica, MA). 48h after transfection, 786-O, Caki-1 cells, or HK2 cells were seeded in XF96-well plates (20,000 cells per well in 80 μL medium). The XF96 sensor cartridge was hydrated with 200 μL calibration buffer per well overnight at 37 °C. For OCR measurement, medium was switched to RPMI (pH 7.4, serum free, 11.10 mM D-glucose, 2.05 mM glutamine) after 12 h, followed by incubation for 1 hour at 37 °C without CO_2_. For ECAR measurement, 2.05 mM glutamine was added into the medium. The sensor cartridge was loaded with assay media (ports A, B and C), oligomycin (1 μM final concentration, port A), carbonyl cyanide 4-(trifluoromethoxy) phenylhydrazone (FCCP, 0.5 μM final concentration, port B) and rotenone and antimycin A (each at 1 μM final concentration, port C) to perform the OCR measurement. Glucose (10 mM final concentration, port A), oligomycin (1 μM final concentration, port B) and 2-deoxy glucose (50 mM final concentration, port C) were added for ECAR measurement. The calibration plate was loaded to equilibrate the sensor cartridge and the standard protocol was started. OCR and ECAR value between different groups was normalized to cell numbers.

#### In vivo mouse experiments

Animal experiments were performed in accordance with the Guiding Principles in the Care and Use of Animals (China) and were approved by the Laboratory Animal Ethics Committee of Xiamen University. Cells (2 x 10^7 786-O or Caki-1) mixed with Matrigel (BD Biosciences, San Jose, CA) in a final volume of 0.1 mL were injected subcutaneously in Balb/c nude mice. Mice were randomized into 2 groups of 5 mice/group 20 days after cell injection. Tumor bearing mice were injected i.p. with 75 mg/kg tozasertib every 48 hours for two weeks. Tumor sizes and mouse body weights were measured periodically.

#### Statistical analysis

R (v.4.1.3) software was used to perform the statistical analyses. The differences in the NDC80 expression levels between normal tissues and tumors in ccRCC cohorts were determined with the Wilcoxon Signed-Rank test. The Kruskal-Wallis test was used to assess differences in hub gene expression in patient samples stratified by clinical parameters (Fig. S4C-F). The Kolmogorov-Smirnov test and logistic regression were used to analyze the relationship between NDC80 expression and clinicopathologic parameters. The Spearman method was used for the correlation of NDC80 expression with the expression of other genes and immune cells. Data from IHC, immunoblotting, cell migration, invasion, cell cycle distribution and mitochondrial function were analyzed by GraphPad Prism 8 using t test.

