## Supplemental Figures for "NDC80 status pinpoints mitotic kinase inhibitors as emerging therapeutic options in clear cell renal cell carcinoma"

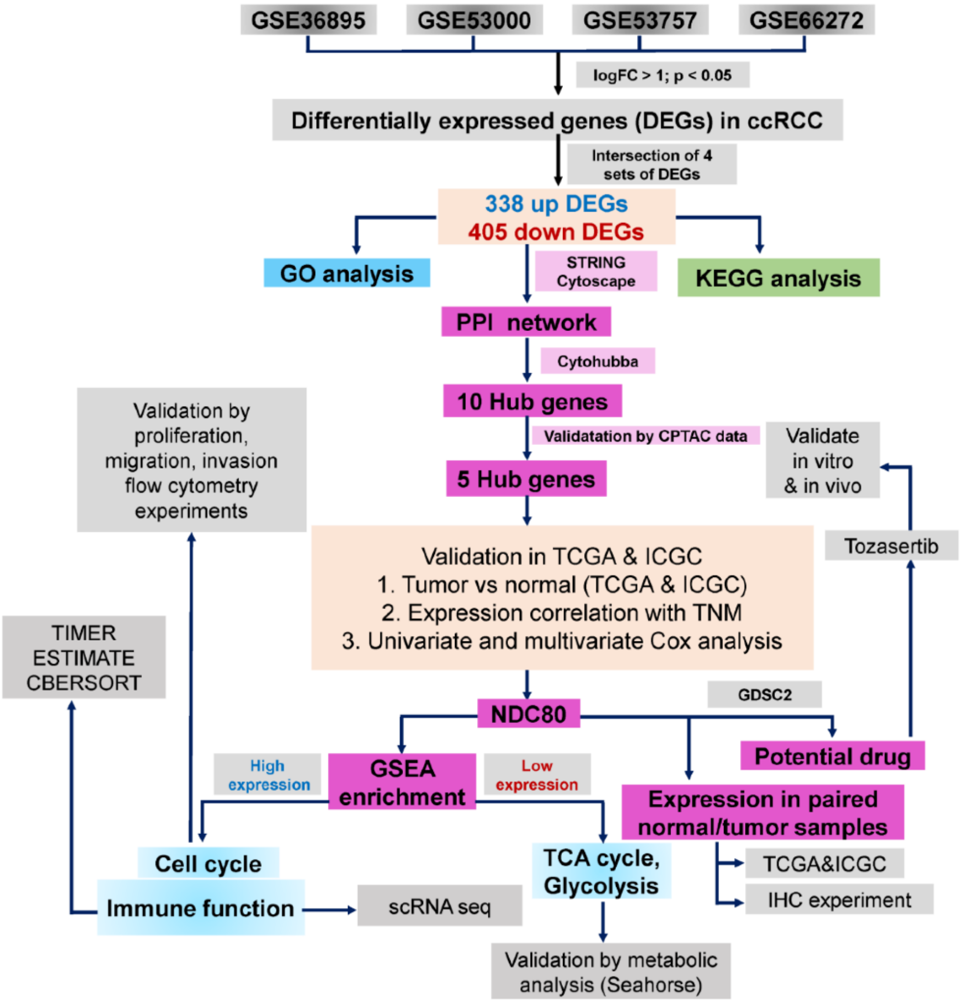


**Figure S1 Schematic of the various datasets and algorithms used to identify NDC80 as a prognostic marker and validation experiments**

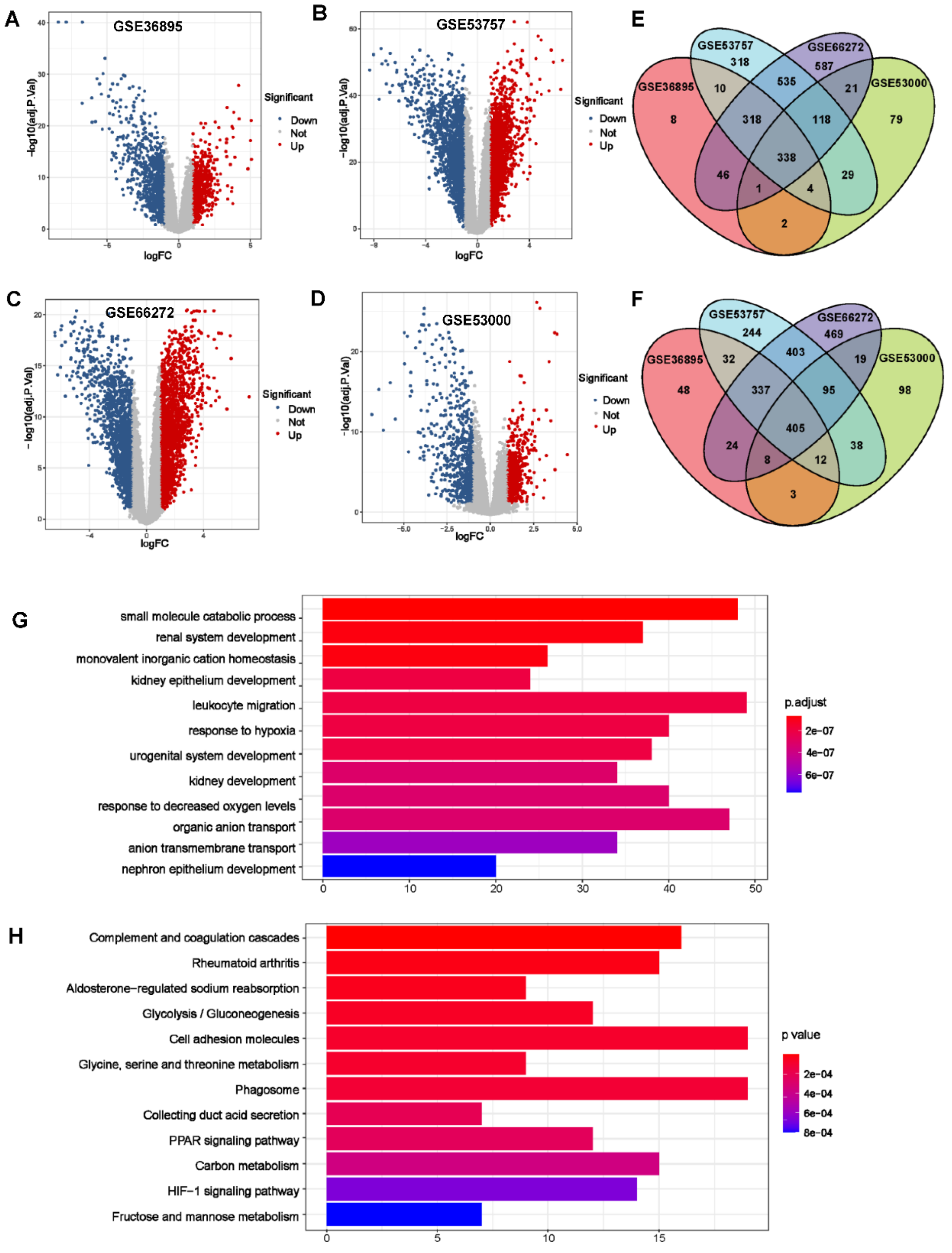


**Figure S2 Functional classification of differentially expressed genes (DEGs) of four ccRCC mRNA expression datasets in GEO**

**(A-D)** Volcano plot of DEGs identified in GSE36895, GSE53757, GSE66272 and GSE53000, respectively. Blue and red dots represent significantly downregulated and upregulated genes.

**(E, F)** Venn diagram of upregulated (E) and downregulated (F) DEGs in the four GEO cohorts.

**(G, H)** Biological Process (BP) Gene Ontology terms (G) and KEGG pathways (H) enriched in the sets of 338 upregulated and 405 downregulated DEGs common to all four ccRCC cohorts.

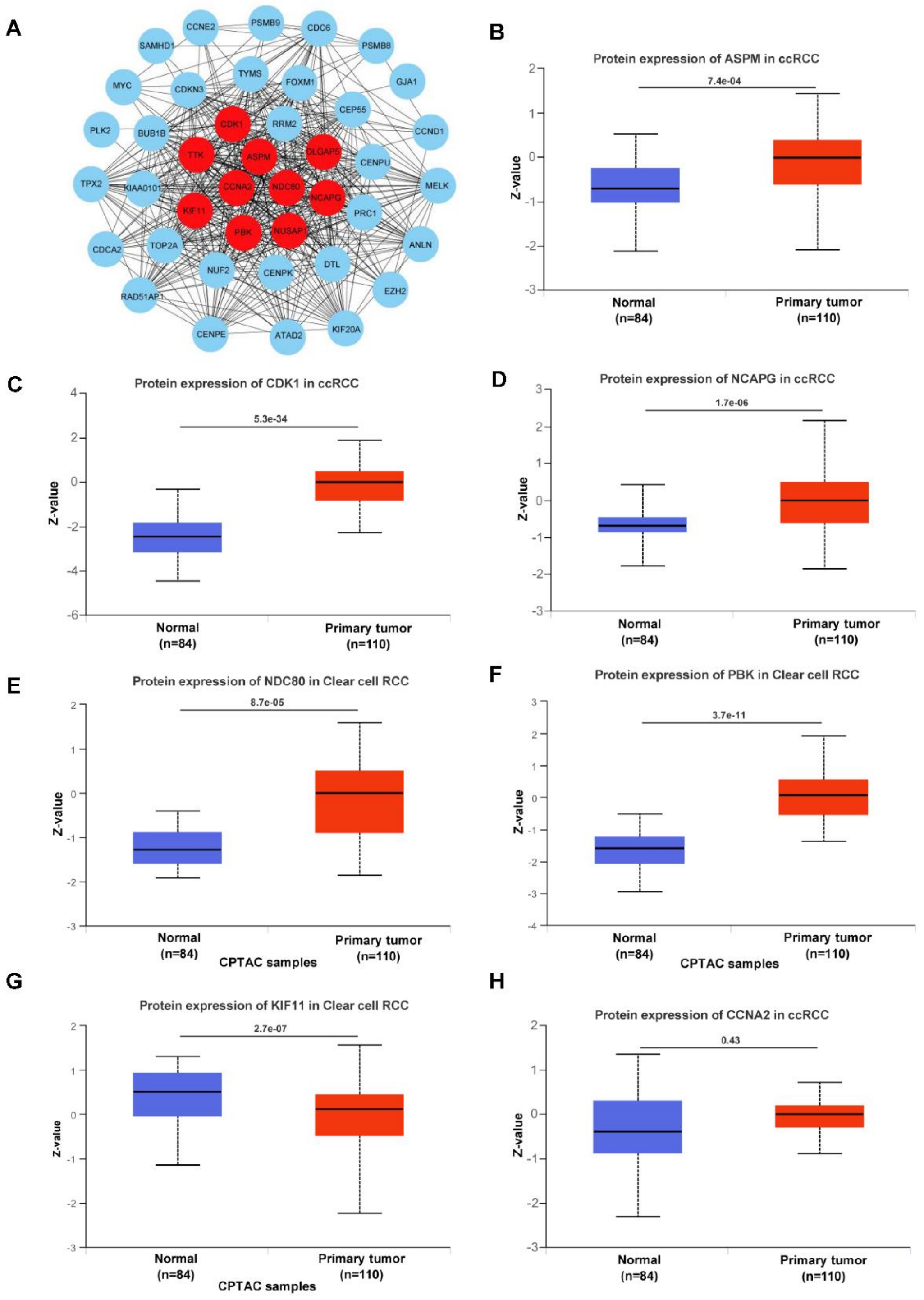


**Figure S3 Protein-protein interaction network and protein expression level of hub genes in ccRCC patients**

(**A**) Protein-protein interaction network of DEGs. Ten hub genes in DEGs were labeled with red.

(**B-H**) Protein expression level of seven hub genes in ccRCC patient samples based on UALCAN CPTAC database.

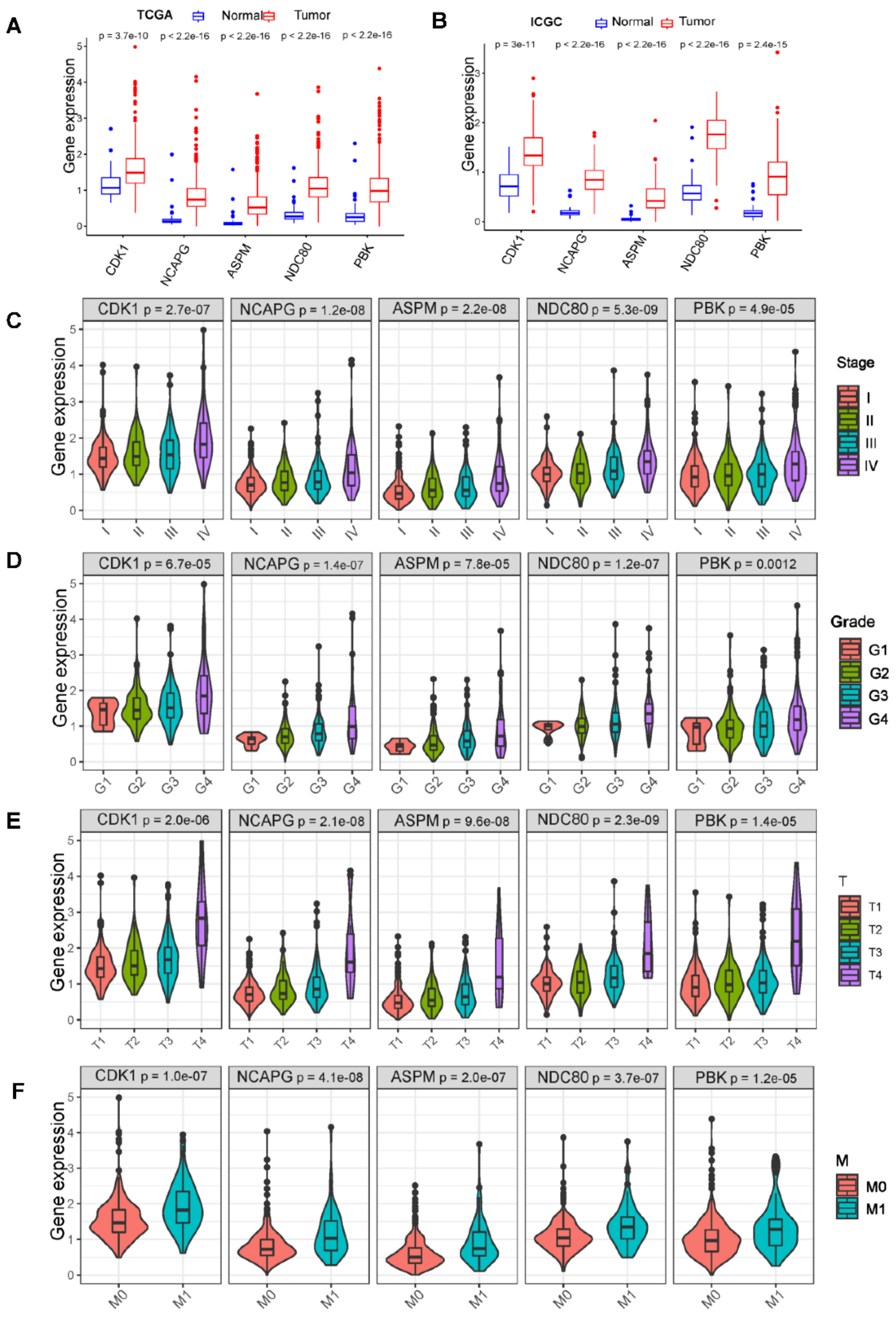


**Figure S4 Expression of CDK1, NCAPG, ASPM, NDC80 and PBK in ccRCC patients**

**(A, B)** Expression of five hub genes in normal and ccRCCs based on TCGA and ICGC database, respectively.

**(C-F)** Distribution of CDK1, NCAPG, ASPM, NDC80 and PBK expression in the stage classification, grade classification, T classification and M classification. P values were determined with the non-parametric Kruskal-Wallis test, except in (F) where the Wilcoxon test was used.

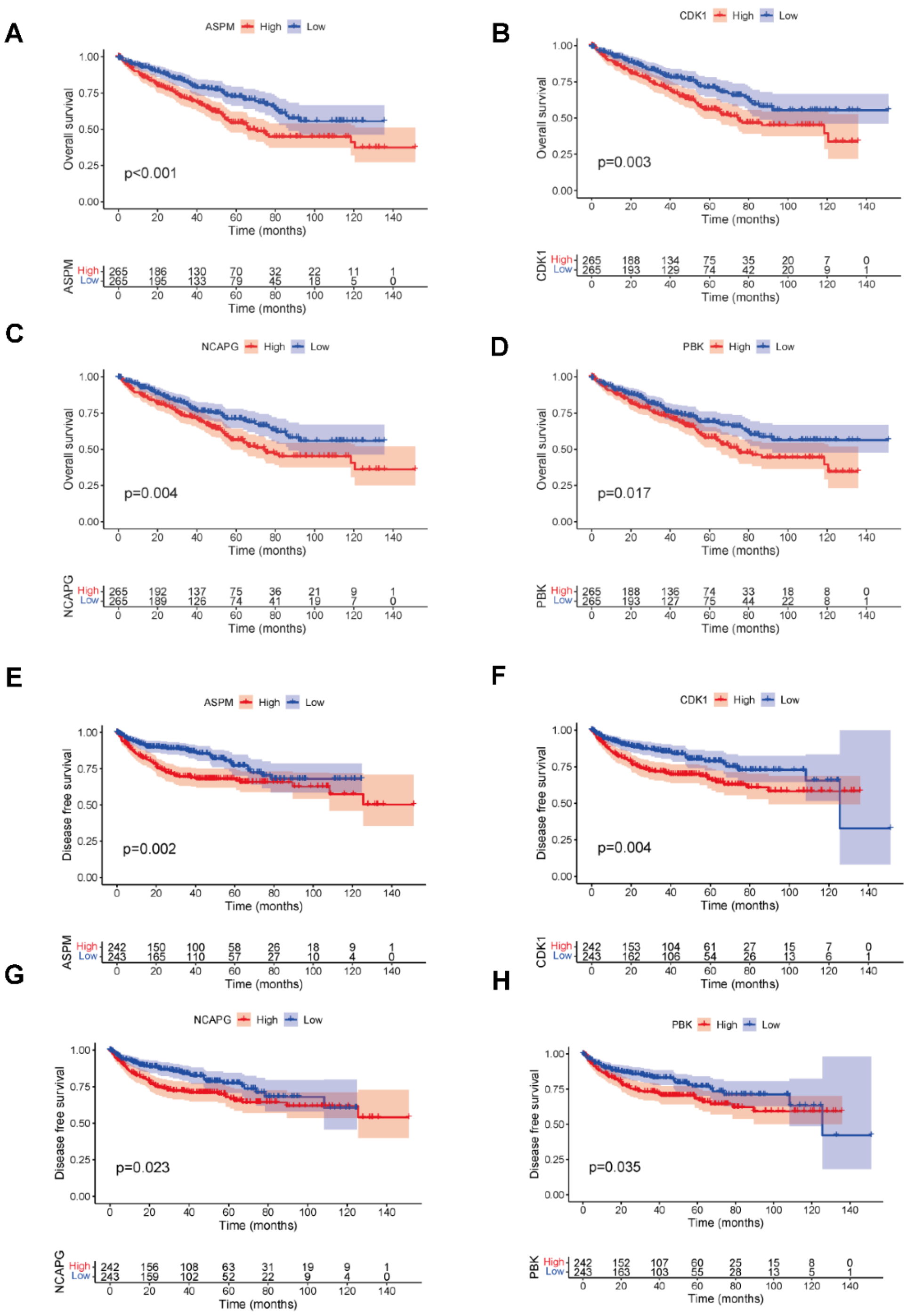


**Figure S5 Kaplan–Meier plots of overall survival (OS) and disease-free survival (DFS) of ccRCC patients based on the expression of hub genes**

**(A-D)** OS ccRCC patients based on the expression of ASPM, CDK1, NACAPG and PBK. Patients were stratified into high and low expressing groups based on the median expression level of the respective mRNAs.

**(E-H)** DFS ccRCC patients based on the expression of ASPM, CDK1, NACAPG and PBK. Patients were stratified into high and low expressing groups based on the median expression level of the respective mRNAs.

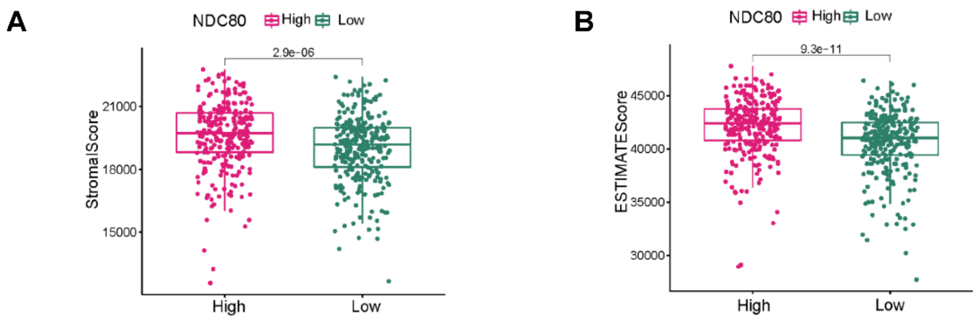


**Figure S6 Stromal and ESTIMATE scores depending on NDC80 expression**

**(A, B)** ESTIMATE algorithm was utilized to calculate the Stromal and ESTIMATE scores between low and high expression of NDC80 cohorts in TCGA database, respectively.

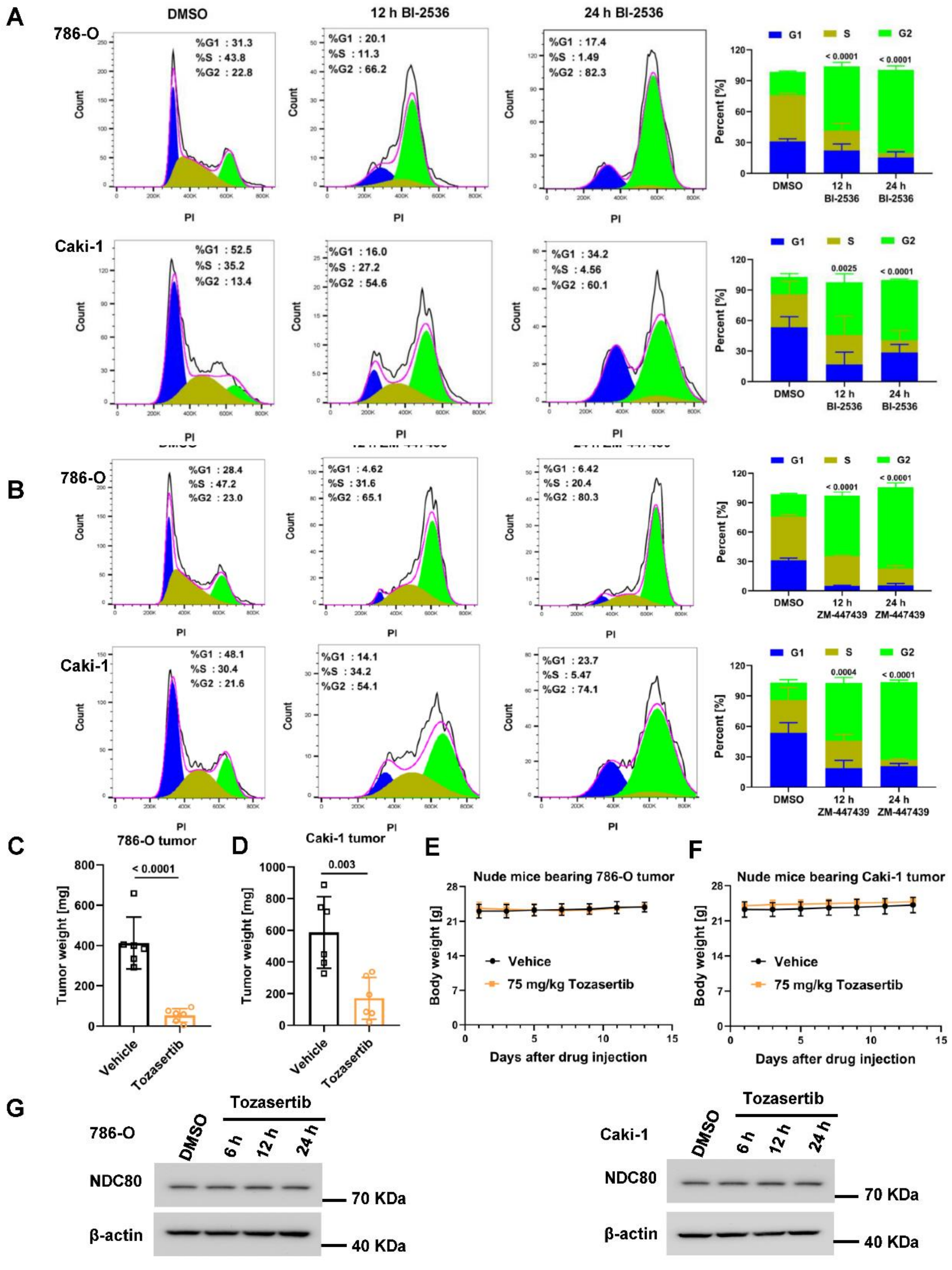


**Figure S7 Effect of mitotic kinase inhibitors on cell cycle progression, tumor weights and body weights of tumor bearing BALB/c nude mice**

**(A, B)** Flow cytometry was performed to detect the cell cycle after treated with DMSO or 12- or 24-hours BI-2536 and ZM447439 under relative IC50 dose in 786-O and Caki-1 cells, respectively (n = 3, averages +/- standard deviation).

**(C, D)** Weights of 786-O or Caki-1 tumors after treatment with 75mg/kg tozasertib or vehicle for two weeks (n = 6, averages +/- standard deviation).

**(E, F)** Body weights of tumor bearing mice in (C and D); n = 6, averages +/- standard deviation.

(**G**) 786-O or Caki-1 cells were treated with 1.34 μM (IC50 dose) and 3.73 μM (IC50 dose) tozasertib respectively for the indicated times, and NDC80 levels were determined by immunoblotting.
